# Environmental DNA Barcode Sequence Capture: Targeted, PCR-free Sequence Capture for Biodiversity Analysis from Bulk Environmental Samples

**DOI:** 10.1101/087437

**Authors:** Shadi Shokralla, Joel F. Gibson, Ian King, Donald J. Baird, Daniel H. Janzen, Winnie Hallwachs, Mehrdad Hajibabaei

## Abstract

Environmental DNA analysis using PCR amplified marker genes has been a key application of high-throughput sequencing (HTS). However, PCR bias is a major drawback to gain accurate qualitative and quantitative biodiversity data. We developed a PCR-free approach using enrichment baits for species-specific mitochondrial cytochrome c oxidase 1(COI) DNA barcodes. The sequence capture was tested on species-rich bulk terrestrial and aquatic benthic samples. Hybridization capture recovered an average of 6 and 4.7 more arthropod orders than amplicon sequencing for terrestrial and benthic samples, respectively. For the terrestrial sample, the four most abundant arthropod orders comprised 94.0% of the sample biomass. These same four orders comprised 95.5% and 97.5% of the COI sequences recovered by amplification and capture, respectively. Hybridization capture recovered three arthropod orders that were detected by biomass analysis, but not by amplicon sequencing and two other insect orders that were not detected by either biomass or amplicon methods. These results indicate the advantage of using sequence capture for a more accurate analysis of biodiversity in bulk environmental samples. The protocol can be easily customized to other DNA barcode markers or gene regions of interest for a wide range of taxa or for a specific target group.

## Introduction

The application of DNA sequence information for the identification of living organisms (e.g., DNA barcoding) has revolutionized biodiversity science (1,2). Customized, public databases of DNA barcodes have been assembled through concerted efforts such as the International Barcode of Life (iBOL) project. These databases employ standardized gene regions for different groups of organisms. For example, cytochrome *c* oxidase subunit I (*COI*) has been established as the standard DNA barcode for animals (1). An increasing number of animal species have been DNA barcoded and their sequences are available publicly through the GenBank and BOLD online data portals (3,4). Similarly, other DNA barcode markers have been established and public libraries built for other kingdoms of life: 16S ribosomal DNA (*16S*) for bacteria and Archaea (5); 18S ribosomal DNA (*18S*) for protists (6,7); the nuclear internal transcribed spacer (*ITS*) for fungi (8); and *rbcL* and *matK* gene regions for plants (9).

Initial DNA barcoding efforts were based on dideoxy chain-termination (“Sanger”) sequencing (10). While scalable, this approach is limited to producing single DNA barcode sequences for one individual at a time. High-throughput sequencing (HTS) (e.g., Illumina MiSeq) has been established as a viable means of assessing DNA barcode-based biodiversity from individuals (11-13) and mixed tissue samples (14-16). The HTS approach can produce a large number of sequences at a greatly reduced overall cost (11-13,15,17). All of these previous examples of massively parallel recovery of DNA barcode gene(s) have relied upon PCR amplification prior to HTS.

Protein-coding genes, such as *COI*, display more sequence variation between individuals and species as compared to ribosomal markers (18). Hence, they are generally considered more discriminating for the purposes of species identification and biodiversity analysis. However, any oligonucleotide primers designed for amplification of protein-coding gene regions may require the addition of degenerate or inert sites to improve taxonomic coverage. The use of degenerate PCR primer sets can still lead to amplification bias, especially when utilized to amplify environmental mixtures with high sequence diversity. Meanwhile, this bias can be exaggerated by the exponential nature of PCR amplification throughout the PCR cycles. This phenomenon can lead to over-amplification of some DNA templates at the expense of non- or under-amplification of other templates. Another possible effect of PCR amplification is the erroneous recovery of nuclear copies of mitochondrial DNA (i.e., NUMTs or pseudogenes). This amplification, sequencing, and reporting of paralogous gene regions can reach substantial rates in some instances (11,19).

It is important to note that primer amplification bias is not due solely to degeneracy and primer mismatch. Primer amplification bias exists even when highly conserved primers are used on mixed tissues. This bias is likely due to differences in GC content of amplicons, amplicon length, annealing temperature, overall genetic diversity of the initial mixture, and the binding energies of the primers themselves (20-22). In order to account for PCR bias, most HTS studies design primers to amplify taxonomic groups of interest. For example, Zeale et al. (23) designed *COI* primers to amplify 12 orders of insects known to make up the diet of bats prior to amplifying and sequencing bat gut contents. In addition to taxon-specific primer design, the employment of multiple PCR reactions with different primer sets has been shown to lessen the impact of primer bias in HTS research. The multiple primer set approach has been especially effective when attempting to recover sequence data from a broad taxonomic range. In mixtures of arthropod tissue from both freshwater benthos (24) and terrestrial Malaise samples (15), the use of multiple primer sets recovers a greater proportion of taxa present than any single primer set alone.

In addition to the possibility of PCR primer bias producing false negative results due to selective exclusion of some species present in a mixture, the issue of organismal abundance has been raised. With the possibility of some DNA templates being selectively amplified over others during the PCR amplification process, relative abundance of DNA sequences cannot be used as a proxy for organismal abundance. Some have attempted to reconcile this discrepancy through the estimation of DNA barcode copy numbers, but this must be calculated one species at a time under controlled lab conditions (25).

The necessity of shorter target gene regions suitable for HTS has resulted in a move towards the use of shorter regions of *COI* contained wholly within the standard *COI* barcode region (14,15, 26,27). In each case, these shorter gene regions are tested *in silico* to ensure that the shorter region can still provide adequate species-level discrimination. In contrast, similar tests of species-level discrimination for other potential marker sequence regions (especially non-protein-coding loci) have given mixed results. For example, short segments of the *ITS* region have proven to be valuable for identification in fungi (28). On the other hand, the nuclear 16S/18S SSU rDNA region has been shown to lump species together producing artificially low reports of biodiversity (29,30).

Hybridization capture followed by HTS has been used in previous instances, usually focusing on individual test species. For instance, the capture method has been used to investigate vertebrate/virus genomic integration (31-33), the phylogeography of small sets of vertebrate species (34,35), and ancient plant DNA (36). When used to recover DNA regions associated with human genetic disease, the capture method was able to supply accurate copy number information for multiple gene regions (37). Also using humans as test organisms, capture protocols were used to recover and reassemble complete individual mitochondrial genomes from mixed starting material (38).

We propose the use of hybridization capture methods, short DNA barcode regions, and HTS to eliminate the effects of PCR bias on biodiversity assessments of mixed environmental samples. We hypothesize that this approach can improve recovery of DNA barcode data from different types of bulk environmental samples as compared to PCR-based analysis. Hence, a capture-based approach should report fewer taxonomic false negatives as compared to a PCR-based method. Also, relative sequence abundance data should be more detailed and accurate than PCR-based relative sequence abundance data. This hypothesis is tested using capture baits designed from a DNA library representing 26 orders of Arthropoda and the tested tissue samples collected via two, distinct arthropod-targeting sampling methods. Malaise traps typically capture terrestrial insects, whereas benthic sampling using a pond net recovers a broad range of aquatic arthropods.

## Material And Methods

### Field sampling

A Malaise sample was collected at Bosque Humedo, Area de Conservación Guanacaste, northwestern Costa Rica (latitude 10.85145°N; longitude −85.60801°W; altitude 290 m; date January 24–31, 2011). The sample was collected directly into 95% ethanol, and frozen at −20°C until thawed and processed. In addition, three benthic samples (A, B, and C) were collected from Wood Buffalo National Park in Alberta, Canada (58.60273°N; 111.52612°W) in June, 2011. The samples were collected 50 meters apart from each other and were taken from the edge of the emergent vegetation zone into the submerged vegetation zone at each site. Samples were collected using a standard pond net with a sterile 400µm mesh net and attached collecting cup attached to a pole. Effort was standardized at 2 minutes per sample. Sampling was conducted by moving the net up and down through the vegetation in a sinusoidal pattern while maintaining constant forward motion. Samples were preserved in 95% ethanol in the field, and kept cold until processed. All DNA-friendly best practices—including wearing clean gloves to collect and handle samples in the field and laboratory and decontaminating nets between samples—were followed to minimize the risk of DNA contamination between sites.

### Biomass calculation

For the Malaise sample, all 1,066 morphologically identifiable individuals were isolated, identified to order morphologically, photographed, measured for total body length (excluding antennae) and maximum thoracic width, and tissue subsampled for individual DNA extraction. Dry biomass of each individual was estimated using order-specific length-width formulae (39).

### DNA extraction

The Malaise sample and each of the benthic samples were individually blended in 95% ethanol and the resultant slurry was transferred to multiple sterile 50 mL Falcon tubes. After ethanol evaporation of the slurry at 56°C, the dried mixture was divided into three lysing matrix tubes “A” (about 100 mg each) and homogenized using the MP FastPrep-24 Instrument (MP Biomedicals Inc.) at speed 6 for 40 sec. Total DNA of this homogenized slurry was extracted using the Nucleospin tissue kit (Macherey-Nagel Inc.) following the manufacturer’s instructions and eluted in 50 µL of molecular biology grade water.

### Amplification-based approach

For all samples, two fragments within the standard *COI* DNA barcode region were amplified with two primer sets (F230R and BR5 for the Malaise sample; AD and BE for the three benthic samples), in a two-step PCR amplification regime (14). The primer sequences are as follows, F:

5`.GGTCAACAAATCATAAAGATATTGG.3` (40), 230_R: 5`.

CTTATRTTRTTTATICGIGGRAAIGC.3` (this paper), A_F: 5`.

GGIGGITTTGGIAATTGAYTIGTICC.3`, B_F:5`.CCIGAYATRGCI-TTYCCICG.3`, D_R:

5`.CCTARIATIGAIGARAYICCIGC.3` (24); and R5: 5`.GTRATIGCICCIG-CIARIAC.3` (15). The first PCR used *COI*-specific primers and the second PCR involved Illumina-tailed primers. The PCR reactions were assembled in 25 µL volumes. Each reaction contained 2 µL DNA template, 17.5 µL molecular biology grade water, 2.5 µL 10× reaction buffer (200 mM Tris–HCl, 500 mM KCl, pH 8.4), 1 µL MgCl_2_ (50 mM), 0.5 µL dNTPs mix (10 mM), 0.5 µL forward primer (10 mM), 0.5 µL reverse primer (10 mM), and 0.5 µL Invitrogen’s Platinum *Taq* polymerase (5 U/µL). The PCR conditions were initiated with heated lid at 95°C for 5 min., followed by a total of 30 cycles of 94°C for 40 s., 46°C (for both primer sets) for 1 min., and 72°C for 30 s., and a final extension at 72°C for 5 min., and hold at 4°C. Amplicons from each sample were purified using Qiagen’s MiniElute PCR purification columns and eluted in 30 µL molecular biology grade water. The purified amplicons from the first PCR were used as templates in the second PCR with the same amplification condition used in the first PCR with the exception of using Illumina-tailed primers in a 15-cycle amplification regime. All PCRs were done using Eppendorf Mastercycler ep gradient S thermal cyclers and negative control reactions (no DNA template) were included in all experiments. PCR products were visualized on a 1.5% agarose gel to check the amplification success.

### Selecting capture targets and probes design

The most important factor in designing a successful Sequence Capture Developer experiment is the quality of the input sequence used to select the target and capture probes. Conservative, hypervariable regions and copy number variation can introduce problems at the design stage (31,35). These problems increase significantly when creating designs for multiple species where some of the taxa are less represented in the database. In this study, a total of 79,215 *COI* DNA barcodes (47,631,471 bp) representing 26 arthropod orders, 559 families, and 4,035 genera were downloaded from both GenBank and BOLD in October, 2012. The 98% similarity clustering of the downloaded sequences resulted in 46,762 unique clusters (28,365,304 bp) (Supplementary Table S1). Sequence Capture Developer probes were designed with the help of the NimbleGen probes design team. A total of 2.1 million 50-105 mer probes (baits) were designed to cover all target clusters. Uniformity of probe abundance was considered, especially for highly conservative regions. The baits design covers 26,918,673 target bases (94.9%) from 46,762 (100%) of the clusters. Four Sequence Capture Developer probes reactions were ordered from NimbleGen, catalog number 06740278001.

### Library preparation for Illumina MiSeq sequencing

For each of the Malaise and benthic samples, a total of 3 µg total DNA of each sample was sheared using a Covaris S220 Focused ultra-sonicator, resulting in a range of 300-800 bp fragments of each sample. The fragmentation efficiency was tested by running the fragmented DNA on an Agilent Bioanalyzer and DNA 1000 chip. Duplicate libraries with unique indexes were prepared with KAPA LTP library preparation kit (Catalog no. KK8230) according to the manufacturer’s protocol.

### Pre-capture pooling and hybridization

For the three benthic samples, the indexed libraries were quantified and equimolar concentrations of each were pooled before hybridization. According to the NimbleGen SeqCap EZ SR user guide v4.2, the pooled and un-pooled indexed libraries were blocked with a blend of COT DNA (Sigma Aldrich, catalog number: 05480647001), MB-grade fish sperm DNA (Roche Diagnostics, catalog number: 11467140001), and plant capture enhancer as a part of Sequence Capture Developer Reagent (Roche Diagnostics, catalog number: 06684335001) in addition to the universal sequencing adaptors and the used indexes. The blocked indexed libraries were hybridized to aliquots of SeqCap EZ library at 47°C for 72 hours with the thermocycler’s head lid maintained at 57°C. The captured libraries were recovered using Streptavidin Dynabeads and washed twice with 1x stringent wash buffer preheated to 47°C followed by subsequent washes with wash buffer I, II, and III and finally recovered with 50 µL of molecular biology grade water. The captured libraries were amplified, purified, and quantified to be ready for sequencing.

### Illumina MiSeq sequencing

The generated amplicons and captured libraries were sequenced in two MiSeq flow cells (one for amplicons and one for captured libraries) using a V3 MiSeq sequencing kit (300 × 2)(FC-131-1002 and MS-102-3001).

### Bioinformatic processing

For the three benthic samples, a total of 24.4 million sequences were generated from PCR amplification and capture libraries, while a total of 11.0 million sequences were generated from PCR amplification and capture libraries of the Malaise sample. For each sample, the forward and reverse raw sequences were kept un-merged as the size of the capture libraries could be variable. All sequences were filtered for quality using PRINSEQ software (41) with a minimum Phred score of 20, window of 10, step of 5, and a minimum length of 150 bp. A total of 17.6 million sequences for benthic samples (mean: 2.0 million reads/sample) and a total of 9.3 million reads for the Malaise sample (mean: 3.1 million reads/sample) were retained for further processing. USEARCH v6.0.307 (42), with the UCLUST algorithm, was used to de-replicate and cluster the remaining sequences using a 99% sequence similarity cutoff. This was done to de-noise any potential sequencing errors prior to further processing. Chimera filtering was performed using USEARCH with the ‘de novo UCHIME’ algorithm (43). At each step, cluster sizes were retained, singletons were retained, and only putatively non-chimeric reads were retained for further processing. All good quality, non-chimera clusters were identified using the MEGABLAST algorithm (44) against a reference library. This reference library contained all verified *COI* sequences downloaded from the GenBank database January 15, 2015 (N = 883,612 sequences). All MEGABLAST searches were conducted with a minimum alignment length of 100 bp and a minimum similarity of 90% for *COI* taxonomic identification recovery based on unambiguous top matches, and 98% for order-, family-, genus-, and species-level identification recovery. All sequencing data generated has been submitted to Dryad and can be accessed at (link will be added once accepted)

## Results

### PCR-based sequencing

PCR amplification followed by sequencing produced between 285,680 and 2,559,402 total sequences for each sample (Table 1). Following quality filtering between 144,356 (50.5%) and 2,342,372 (91.5%) high-quality sequences remained for analysis. Of these high-quality sequences, between 135,560 (93.9%) and 2,075,555 (88.6%) sequences were identified as *COI* sequences (i.e., had at a minimum of 90% similarity to a reference *COI* sequence in GenBank).

**Table 1.** Summary of number of DNA sequences produced, filtered, and used via four different samples and two different methods.

| Sample | Process | # raw DNA sequences | # sequences passing quality filter | # *COI* sequences | % high quality sequences used in *COI* analysis |
| --- | --- | --- | --- | --- | --- |
| Benthic A | Amplicon | 285,680 | 144,356 | 135,560 | 93.9 |
|  | Capture 1 | 4,331,460 | 3,713,290 | 1,954,225 | 52.4 |
|  | Capture 2 | 4,023,194 | 2,993,002 | 1,939,291 | 64.8 |
| Benthic B | Amplicon | 582,590 | 341,528 | 304,548 | 89.2 |
|  | Capture 1 | 4,042,778 | 2,703,271 | 1,695,684 | 62.7 |
|  | Capture 2 | 3,010,292 | 1,700,512 | 1,001,123 | 58.9 |
| Benthic C | Amplicon | 510,396 | 411,362 | 373,668 | 90.8 |
|  | Capture 1 | 3,438,490 | 2,558,027 | 1,744,542 | 68.2 |
|  | Capture 2 | 4,177,500 | 3,027,892 | 2,046,278 | 67.6 |
| Malaise | Amplicon | 2,559,402 | 2,342,372 | 2,075,555 | 88.6 |
|  | Capture 1 | 4,444,412 | 3,499,795 | 2,278,390 | 65.1 |
|  | Capture 2 | 4,042,698 | 3,436,537 | 2,041,300 | 59.4 |

### Hybridization capture sequencing

Hybridization capture followed by sequencing produced between 3,010,292 and 4,444,412 total sequences for each sample (Table 1). Following quality filtering, between 1,700,512 and 3,713,290 (up to 85.73%) high-quality sequences per sample remained for analysis. Of these sequences, between 1,001,123 and 2,278,390 (up to 68.2%) sequences per sample were identified as *COI* sequences as mentioned before.

### Taxonomic data recovery

For the benthic samples, hybridization capture followed by HTS recovered an average of 18 more orders than PCR amplification followed by HTS. For the Malaise sample, the average increase in order richness was 7. For the benthic samples, hybridization capture followed by HTS recovered an average of 31.8 more families than PCR amplification followed by HTS. For the Malaise sample, the average increase in family richness was 23. For the benthic samples, hybridization capture followed by HTS recovered an average of 37.2 more genera than PCR amplification followed by HTS. For the Malaise sample, the average increase in genus richness was 19.5.

Differences in taxonomic recovery considering only arthropod orders—for which the sampling and capture baits were designed—were also calculated. For the benthic samples, hybridization capture followed by HTS recovered an average of 4.7 more arthropod orders than PCR amplification followed by HTS (Figure 1). For the Malaise sample, the average increase in order richness was 6. For the benthic samples, hybridization capture followed by HTS recovered an average of 15 more arthropod families than PCR amplification followed by HTS. For the Malaise sample, the average increase in arthropod family richness was 24. For the benthic samples, hybridization capture followed by HTS recovered an average of 18.2 more arthropod genera than PCR amplification followed by HTS. For the Malaise sample, the average increase in arthropod genus richness was 21.5.

**Figure 1.**
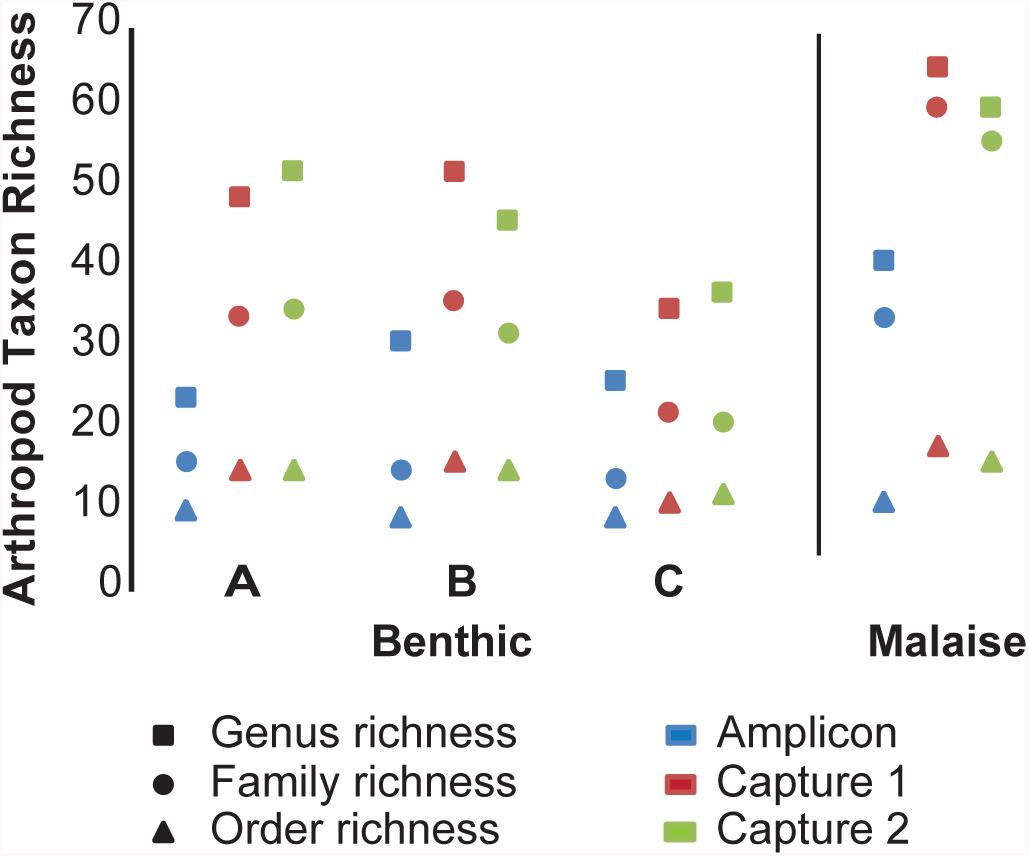
Taxonomic richness recovery at three levels for four different subsamples using PCR amplification and hybridization capture.

### Relative sequence abundance

For the Malaise sample, the four most abundant orders (Diptera, Lepidoptera, Coleoptera, and Hymenoptera) were all arthropods and comprised 94.0% of the biomass of the sample (Supplementary Table S2). These same four orders comprised 95.5%, 97.5%, and 97.4% of the *COI* sequences recovered by PCR amplification, Capture 1, and Capture 2, respectively. Three arthropod orders (Blattodea, Trombidiformes, and Psocoptera) were detected by morphological identification and biomass analysis, but not by any molecular method. Also, Coleoptera was present as a much higher proportion of biomass (35.6%), than as sequence proportion by any of the three methods (amplification 0.08%; capture 0.23% and 0.29%). Hybridization capture recovered three arthropod orders (Collembola, Neuroptera, Trichoptera) that were detected by biomass analysis, but not by PCR amplification. Furthermore, hybridization capture detected two insect orders (Mantodea, Plecoptera), in very low sequence numbers, which were not detected by either biomass analysis or PCR amplification (Supplementary Table S2).

For the Benthic A sample, at the order level, the three most abundant arthropod orders (Diptera, Hemiptera, Podocopida) combined represented 56.0%, 56.4%, and 59.3% of the *COI* sequences recovered for PCR amplification, Capture 1, and Capture 2, respectively. For the Benthic B sample, at the order level, the mentioned three most abundant arthropod orders combined represented 60.8%, 56.2%, and 55.0% of the *COI* sequences recovered for PCR amplification, Capture 1, and Capture 2, respectively. For the Benthic C sample, at the order level, the three most abundant arthropod orders combined composed 93.3%, 92.7%, and 92.9% of the *COI* sequences recovered for PCR amplification, Capture 1, and Capture 2, respectively (Figure 2).

**Figure 2.**
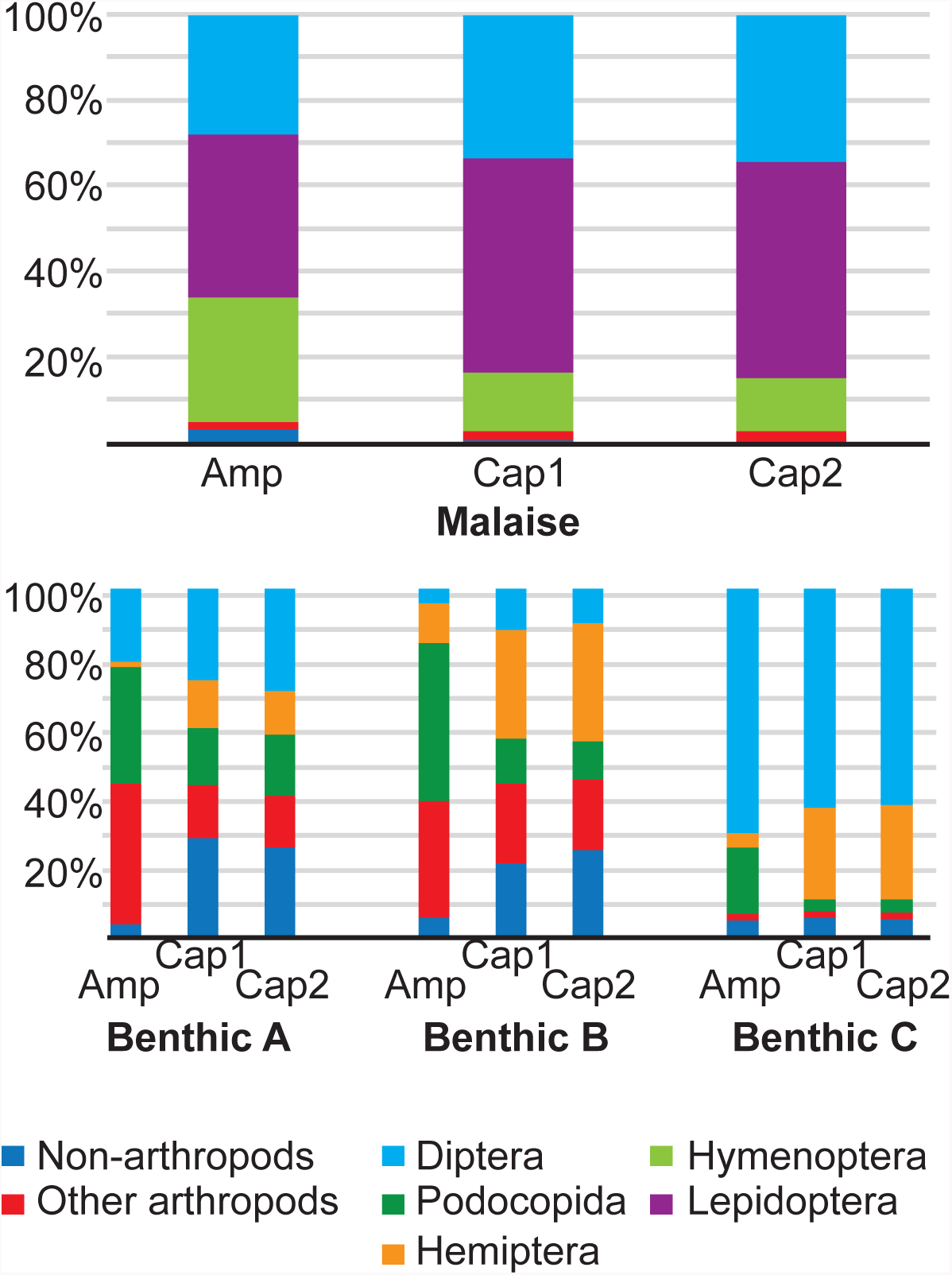
Proportion of DNA sequences identified to order for four different subsamples using PCR amplification and hybridization capture. Amp – sequences recovered via PCR amplification. Cap1, Cap2 - sequences recovered via hybridization capture in the first and second attempts.

A comparison between methods of the proportion of sequences assigned to each arthropod order, family, and genus was used to generate scatter plots and correlation values (Figure 3). Regardless of sample or taxonomic level, the two rounds of hybridization capture produced highly correlated (0.99 to 1.00) proportions assigned to each taxon. The degree of correlation between PCR amplification and hybridization capture varied (0.43 to 0.93), but was consistently below that of hybridization capture round one versus round two. Similarly, Bray-Curtis dissimilarity values calculated from the proportion by taxon matrices at the arthropod order, family, and genus level were calculated (Figure 4). Likewise, dissimilarity values were much lower for hybridization capture round one versus round two comparisons (0.011 to 0.077), than those for PCR amplification versus hybridization capture comparisons (0.166 to 0.615).

**Figure 3.**
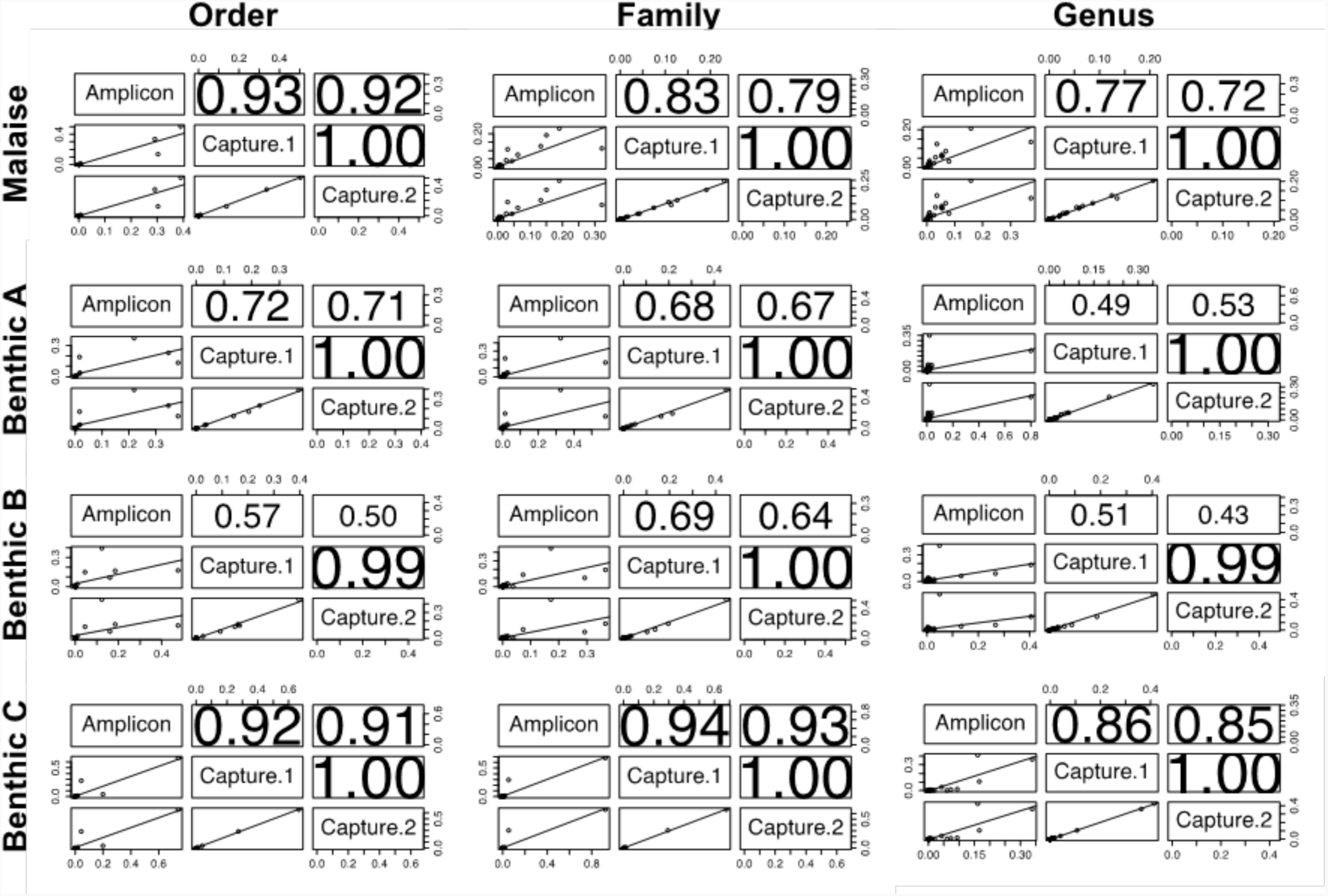
Scatter plots and correlation values of proportion of sequences assigned to each arthropod order, family, and genus for four different subsamples using PCR amplification and hybridization capture.

**Figure 4.**
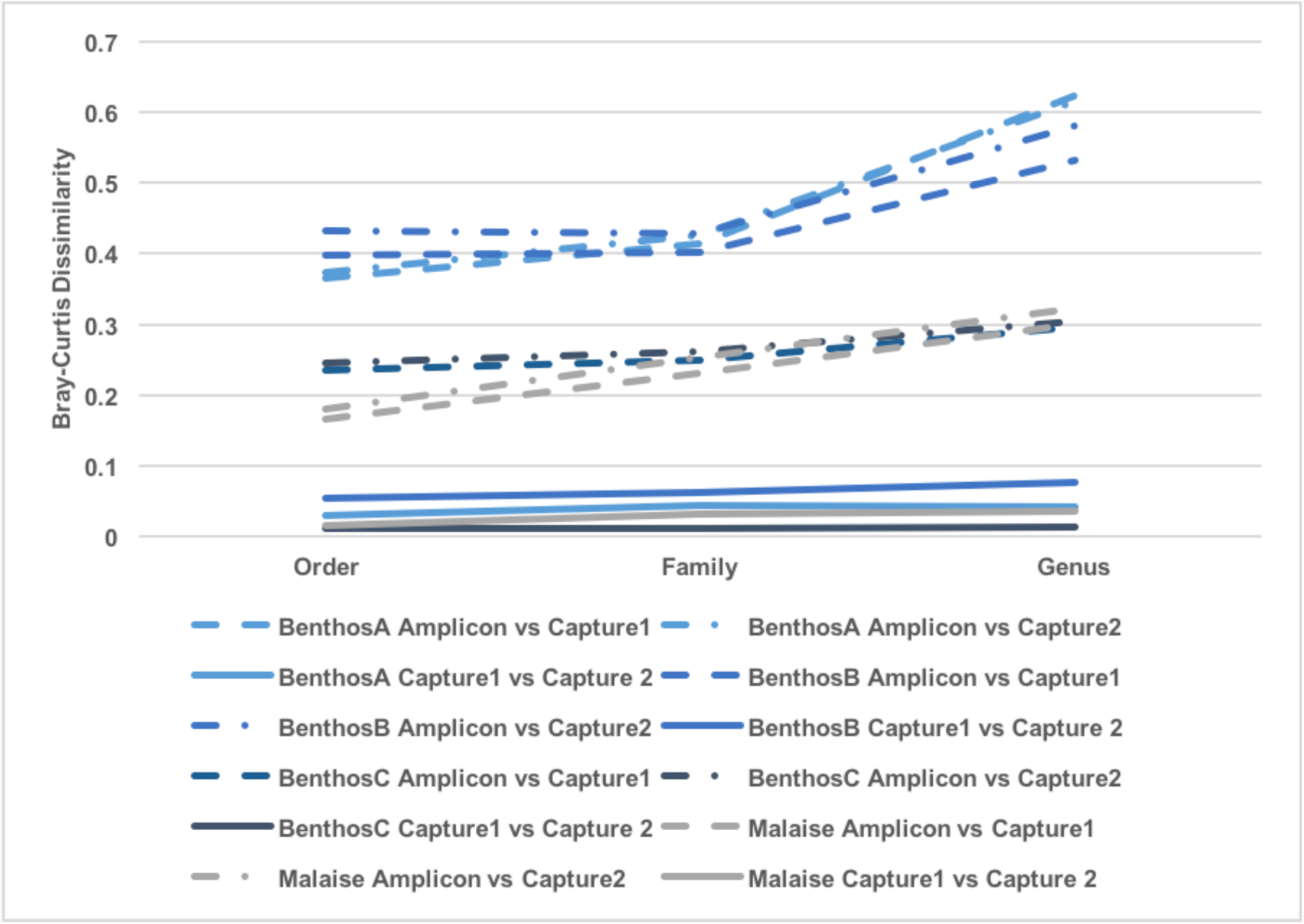
Bray-Curtis dissimilarity values based on proportion of sequences assigned to each arthropod order, family, and genus for four different subsamples using PCR amplification and hybridization capture.

## Discussion

This study is the first targeted attempt at PCR-free DNA barcode recovery from mixed environmental samples. Previous work employed mitochondrial enrichment followed by sequencing of total enriched mtDNA (45). This method required the exclusion of 99.5% of sequence data in order to retain only the 0.5% that comprises informative *COI* DNA barcode data. Liu et al. (46) were able to increase the efficiency of whole mitochondrial genome sequencing to about 42.5% by using a capture microarray followed by HTS in a mock community sample containing equal aliquots of genomic DNA extracts of 49 species. The present method retains 52.4 to 68.2% of the passed filter sequences produced through Illumina MiSeq sequencing for final analysis (Table 1). Whether pooled as three lower diversity samples or retained as one single, high diversity, high sequencing coverage sample, this produces one to two million *COI* sequences per sample for further analysis. This rate of analyzable sequences does not differ between benthic- and Malaise-derived tissue samples.

The recovery of non-target DNA regions has been noted in previous use of capture enrichment in resequencing studies where multiple regions of the genome need to be enriched and analyzed (33,34,37). This occurrence is likely due to the capture of adjacent (“flanking”) DNA regions during the hybridization process. The proportion of recovered sequences not matching targets ranges between 18 and 42% in these previous studies. In the present study, 31.8 to 47.6% of high-quality sequences produced through hybridization capture recovered were not positively identified as *COI*. This rate in target recovery, while similar to previous resequencing studies, differs fundamentally from past mitochondrial enrichment experiments. Previously, hybridization baits were designed and implemented to target a number of different gene regions for a narrow range of organisms (e.g., the entire mitochondrial genome of one species). The present use of hybridization capture targets many versions of one small gene region in order to retain sequence data from a taxonomically wide range of organisms. As such, the rate of non-target sequence generation can be interpreted as either capture of non-*COI* sequences with some affinity for the oligonucleotide baits used, or else sequencing error within the Illumina MiSeq resulting in uninformative sequences. Any truly “flanking” sequences adjacent to the *COI* region targeted could still be retained and identified as *COI* sequences.

Hybridization capture followed by HTS recovered approximately twice as many order, family, and genus names as compared to PCR amplification for the three benthic samples (Table 1, Figure 1). Likewise, hybridization capture followed by HTS recovered an increase in order, family, and genus richness for the Malaise sample. These increases in taxonomic recovery are more pronounced when only the Arthropoda are considered. These increases in biodiversity detection reflect a reduced false negative rate due to the absence of primer amplification bias. The difference in increased taxonomic recovery at each level reveals an important difference between the benthic and Malaise samples. For both Malaise and benthic samples, the majority of the orders recovered only through hybridization capture were non-insects, including other arthropods, vertebrates, diatoms, fungi, and plants. Benthic samples, being aquatic in nature, will include a number of orders not found in a terrestrial sample, especially zooplankton and phytoplankton. To rule out false positives, we considered only the sequencing clusters containing more than ten sequences per cluster. The PCR amplification primers employed were designed for insects and are unlikely to have amplified non-insect orders. The hybridization capture, despite also being designed from insect DNA sequences, employs multiple longer oligos complementary to multiple fragments of the *COI* gene region and was better able to recover non-insect taxa.

It has been stated that PCR primer bias is likely to make conventional barcode copy abundance calculations from HTS environmental DNA studies impossible (e.g., 47). The removal of this bias facilitates a shift towards more meaningful interpretations of sequence numbers recovered from environmental samples and decreases chances of false negatives for taxa with a small biomass or rare taxa (15,17). For all benthic and Malaise samples, there was consistency in relative sequence frequencies between hybridization capture attempts at the arthropod genus, family, and order levels (Figure 2, Table 2). In all samples, PCR amplification recovered a different community profile from hybridization capture (Figures 2, 3; Table 1).

In the Malaise sample, when compared to biomass calculations, the difference in proportion of sequences assigned to each taxon is noticed as an overabundance of Lepidoptera sequences at the expense of Coleoptera. In addition, three orders (Blattodea, Trombidiformes, and Psocoptera) are not detected by HTS methods at all. This result is likely a product of the taxonomic composition of GenBank *COI* libraries. An abundance of tropical Lepidoptera are represented as GenBank records whereas few Costa Rican Blattodea, Psocoptera, and Trombidiformes are included. This lack of likely matches, coupled with our strict 98% similarity cut-off, resulted in a failure to recover these orders. As a further analysis, when similarity cut-offs were loosened to 90% similarity, with the same GenBank library, Blattodea, Trombidiformes, and Psocoptera were recovered with HTS methods, and Coleoptera was recovered as a much greater proportion of sequences (results not shown). However, a relaxed cut-off for sequence analyses such as BLAST could also result in reduced confidence in taxonomic identification of sequences or may result in false positives. As reference DNA barcode libraries are populated with diverse taxa and include better representation of local populations (haplotypes), sequence capture studies can benefit from these additional sequences both for the design of capture probes and in subsequent analyses of sequences. In fact, the use of standard DNA markers (e.g., barcodes) and provisioning reference databases with voucher specimens could facilitate large-scale analyses of environmental samples.

The use of HTS to allow parallel sequencing of multiple genetic markers in both library building and environmental mixture research has been previously proposed and demonstrated (11,12,15). While the present study focused on *COI* region as the most widely used DNA barcodes, the protocol could be easily adapted to include other DNA barcode regions, phylogenetic markers, or functional genes. The presented approach offers the flexibility to be customized to specific target groups and multiple barcoding markers. It also overcomes the challenges in PCR-based methods including primer design in hypervariable markers and the associated bias. The ability to multiplex many samples and recover individual sample data at sufficient sequencing depth is shown here to have no negative effect on biodiversity data recovery. The push to include as many samples and genetic markers as desired in future HTS-based biodiversity studies would be readily accommodated by the current protocol.

## Acknowledgement

We are grateful to Area de Conservación Guanacaste for protecting the forest habitat that we sampled.

## Supplementary Material

**Table S1.**
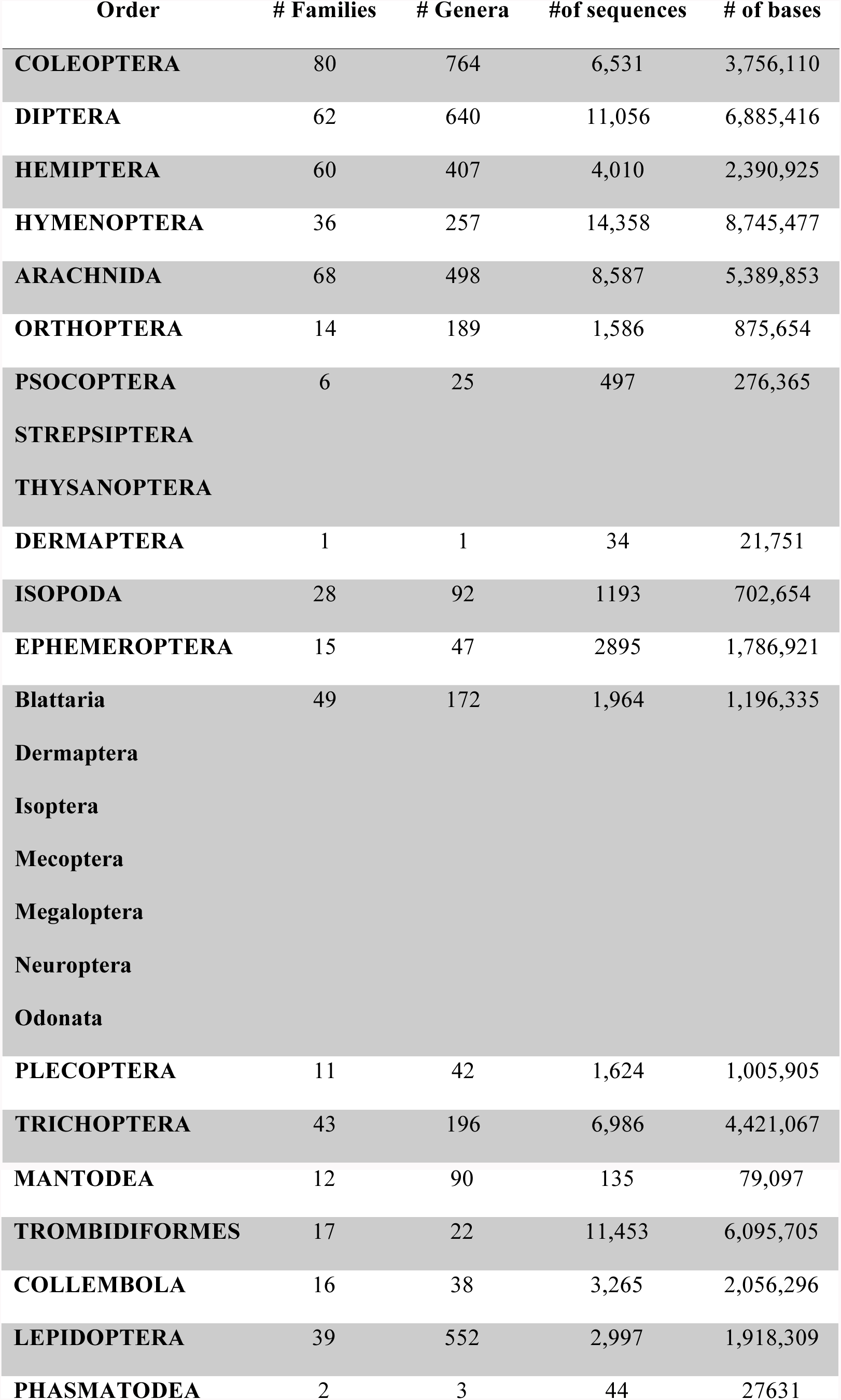
Summary of sequences included in oligo development.

**Table S2.**
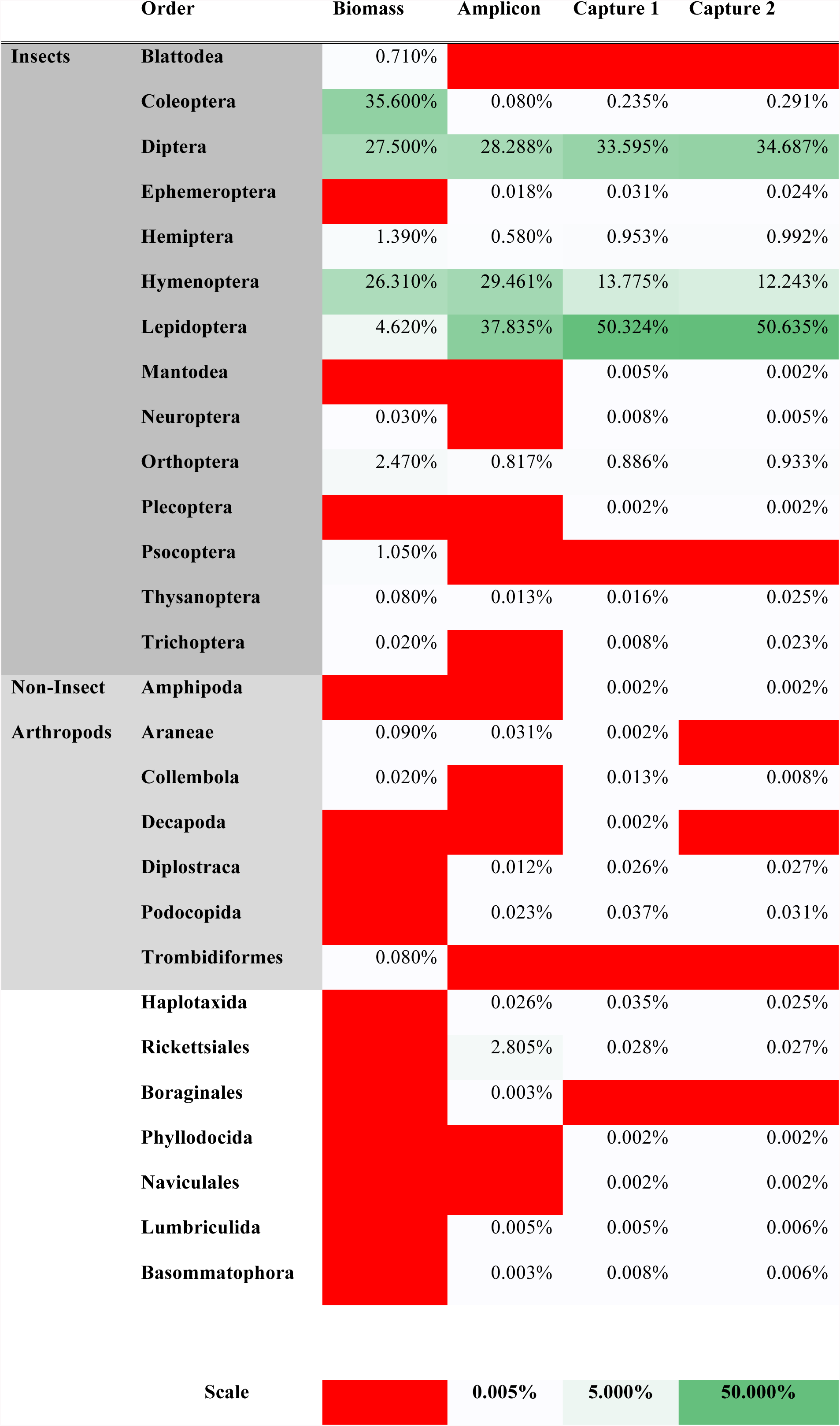
Proportion of calculated total biomass and proportion of *COI* DNA sequences identified to insect, arthropod, and non-arthropod orders via two different methods for a single Malaise sample.

